## Supplementary figures and images for "Phospholipid peroxidation fuels ExoU phospholipase-dependent cell necrosis and supports *Pseudomonas aeruginosa*-driven pathology"

### Figure S1

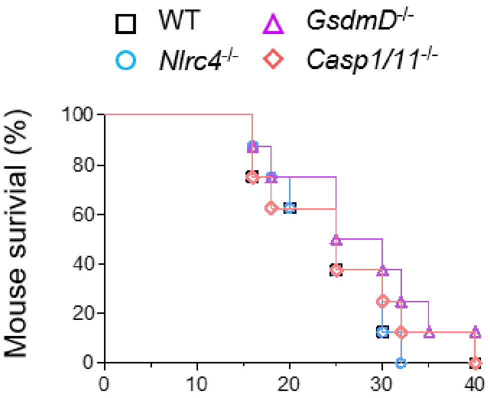

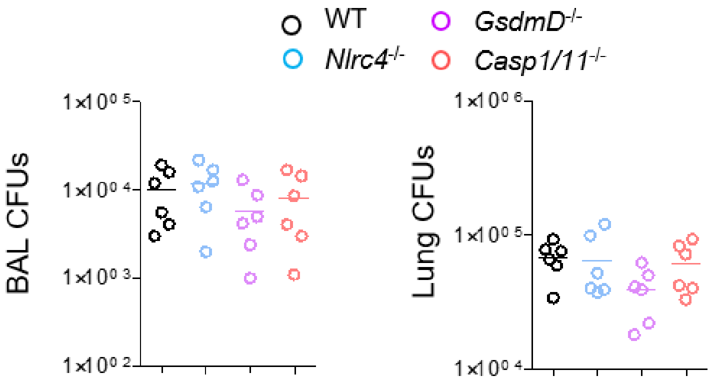

### Figure S2

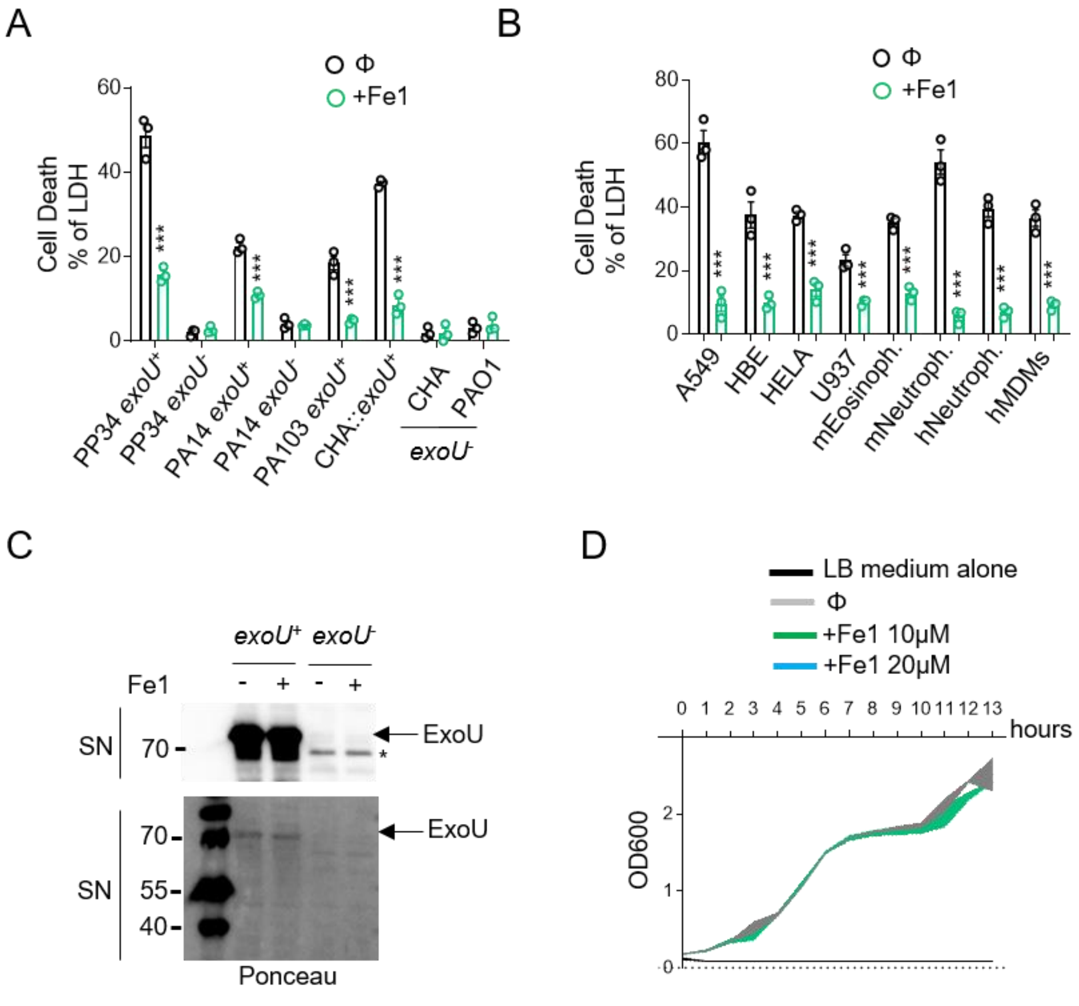

### Figure S3

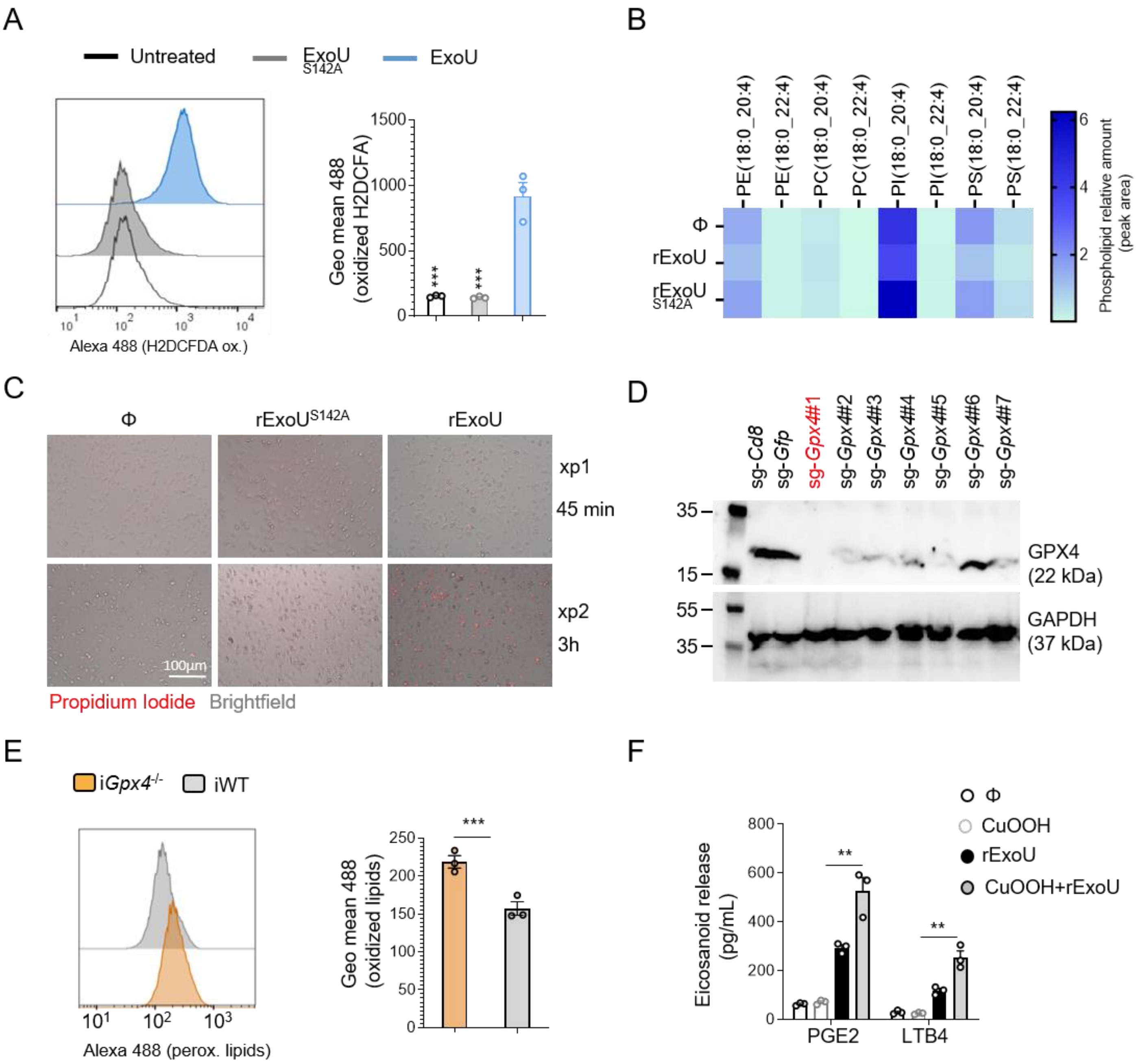

### Figure S4

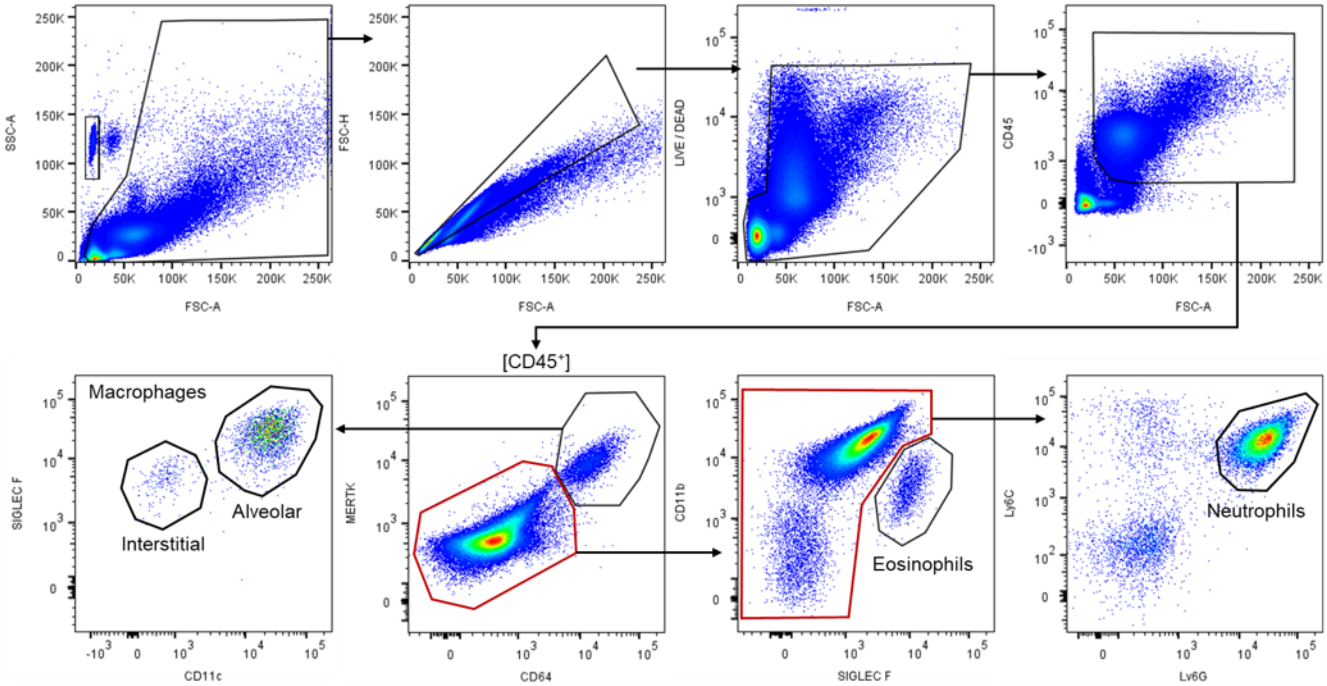

### Graphical abstract

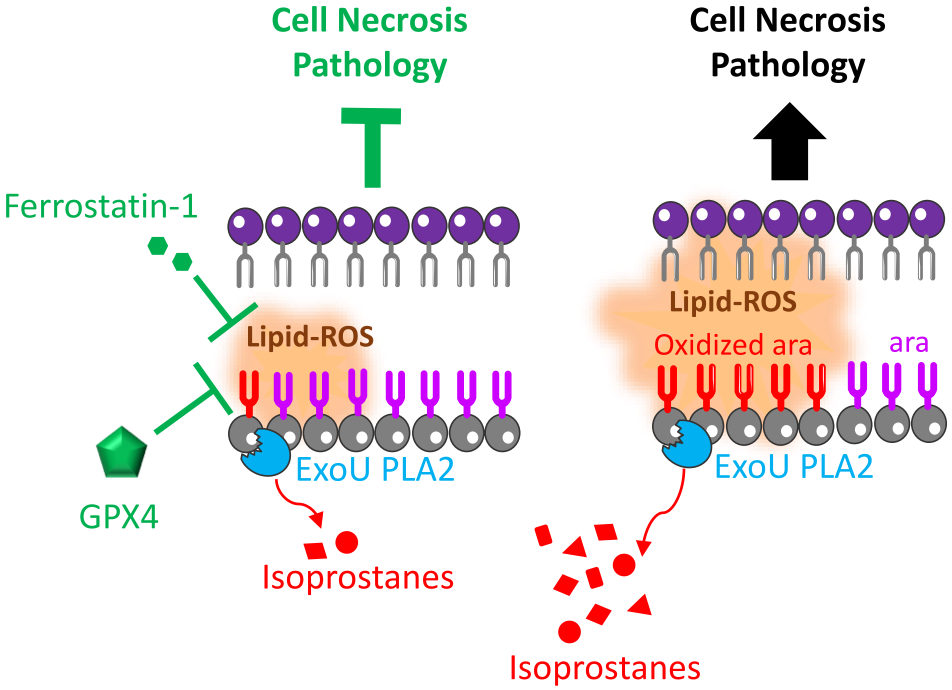
